# Identifying Cross-Cancer Similar Patients via a Semi-Supervised Deep Clustering Approach

**DOI:** 10.1101/2020.11.07.372672

**Authors:** Duygu Ay, Oznur Tastan

## Abstract

The treatment decisions for a cancer patient are typically based on the patient’s diagnosed cancer type. With the characterization of cancer tumors at the molecular level, there have been reports of patients being similar despite being diagnosed with different cancer types. Motivated from these observations, we aim at discovering *cross-cancer* patients, which we define as patients whose tumors are more similar to patient tumors diagnosed with another cancer type. We develop DeepCrossCancer to identify cross-cancer patients that always co-cluster with the other patient from another cancer type. The input to DeepCrossCancer is the transcriptomic profiles of the patient tumors, the age, and sex of the patient. To solve the clustering problem, we use a semi-supervised deep learning-based clustering method in which the clustering task is supervised by cancer type labels and the survival times of the patients. Applying the method to patient data from nine different cancers, we discover 20 cross-cancer patients that consistently co-cluster. By analyzing the predictive genes of the cross-cancer patients and other genomic information available for the patient such as somatic mutations and copy number variations, we identify striking genomic similarities across these patients providing support. The detection of cross-cancer patients opens up possibilities for transferring clinical decisions across patients at a single patient level. The source code is available at github.com/tastanlab/DeepCrossCancer

## 1 Introduction

Cancer cells exhibit numerous genomic alterations compared to normal cells [1] that widely differ across patients that are diagnosed with the same cancer type [2]. This heterogeneity sets significant challenges for designing effective diagnostic and treatment strategies that would work across all patients of a cancer type [3]. The large-scale cancer molecular profiling projects have opened up possibilities for developing more precise therapies [4]. Characterizing the alterations in the cancer cells also enhances the ability to understand the molecular underpinnings of tumor development and progression, which can also inform clinical management. There are two research directions undertaken that rely on the analysis of this rich molecular data that serve these goals.

In the first research direction, molecular subtypes of the same cancer type are sought after to dissect the heterogeneity observed within a cancer type. The ultimate goal is to design treatment regimens tailored for each subgroup and understand what differs them at the molecular level. The identification of breast cancer intrinsic molecular subtypes discovered almost two decades ago by analyzing gene expression profiles of cancer patients is an example of such an approach [5,6]. Similar molecular subtyping efforts have been undertaken for breast cancer with data obtained in the sequencing projects and for other cancer types [5–15].

The second research direction is the pan-cancer analysis of cancer patient-derived molecular data [16]. The aim here is to reclassify human tumor types based on their molecular similarity and get a unified view of multiple types of cancer on commonalities and differences. To this end, the Pan-Cancer Genome Atlas project performed an integrative molecular analysis using multiple types of omic data from 33 different tumor types [17] and using a clustering approach, [18,19] and arrived at 28 distinct molecular subtypes. In other diseases, such as psychiatric disorders, such global cross-disorder analysis has been conducted [20,21].

In this work, we focus on a third strategy, we identify cross-cancer patients. We define a patient as *cross-cancer* patient if they bear molecular similarity to a single or multiple cancer patients in another cancer type. Such similarities have been reported in the literature. A TCGA analysis had revealed that the subtype of breast cancer – the basal-like – bear extensive molecular similarities to high-grade serous ovarian cancer, which has been hard to treat [22]. Similarly, the results on endometrial carcinomas of TCGA demonstrated that 25% of 373 tumors studied that have been classified as high-grade endometriosis by pathologists have molecular similarities to uterine serous carcinomas [23]. In this work, we even look into finer similarities, such as a specific patient being similar to another patient that could be missed in these global analyses.

This approach is different from the subtype discovery efforts because it focuses on similarities not within a cancer type but across cancers. It is also different from the pan-cancer analysis because instead of finding global similarities across a large group of patients, it seeks more patient-specific similarities for a single patient that could be missed in a pan-cancer study due to the small cluster size. The benefit of such an approach is two folds. If there are actionable genomic events, the detection of cross-cancer patients opens up possibilities for transferring clinical decisions from one patient to the other one immediately. Secondly, patients in different cancer types with these unexpected molecular similarities can discover novel cancer-driving mechanisms.

There are two algorithmic contributions to this work. The first contribution is that we define the cross-cancer patient identification problem and propose a novel method to identify cross-cancer patients based on transcriptomic data of tumors biopsied from patients and clinical information. Secondly, we propose a semi-supervised learning algorithm that extends a previous deep learning clustering method designed for subtype discovery [24]. Unlike [24] approach, our model also uses survival information of the patients to guide the clustering along with their diagnosed cancer types. Although the ultimate aim is to reach good clusters of the patients, solving the auxiliary tasks of cancer type classification and the survival prediction serve to learn a good representation of the patients. When we apply this method on patient data from nine cancer, we identify 20 cross-cancer patients. We inspect these patients in the light of other genomic data available such as somatic mutations and copy number variations and find interesting common genomic events.

## 2 Methods

### 2.1 Problem Formulation

Consider a set of *n* patients diagnosed with *m* different cancer types. Let *y_i_* denote the cancer type for the patient *i*, where *y_i_* ∈ {1, …, *m*}. We deem patient *i* a cross-cancer patient if it follows the following conditions: i) There is at least one patient *j* where *i* and *j* always co-clusters over multiple runs of clustering and for which *y_i_* ≠ *y_j_*. ii) Patient *i* is closer to *y_j_* members than its own cancer type *y_i_* members. Note for a pair of similar patients (*i,j*), not necessarily both of the patients will be cross-cancer patients due to the second condition above. While *i* is a cross-cancer patient, *j* might not be a cross-cancer patient and could be a typical member of its own cancer type members. The algorithm proceeds in two steps. The first step involves clustering patients diagnosed with different cancer types. The second step involves clustering multiple times and identifying the patient samples that stably co-clustered with the patient sample(s) from another cancer type to identify the cross-cancer patients. In the next section, we detail these steps. To facilitate the reading of the subsequent parts we provide a summary of notation used in Table S1.

### 2.2 Step 1 - Semi-supervised Deep Clustering

We want to cluster *n* patient tumor samples using the samples’ molecular profiles and additional clinical information. Note that we will use the terms patient and the patient’s tumor sample interchangeably. We denote i-th patient’s feature vector with 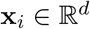. The features are the gene expression data and the patient’s gender and age group. In the clustering step, the m patients presented with the feature vector **x**_*i*_ ∈ *X* are grouped into *k* disjoint clusters, each of which is represented by a centroid **u***_j_*, *j* ≡ 1,…, *k*. **U** will denote the centroid matrix, where *j*-th column is the cluster centroid, **U**_*j*_. We will denote the cluster assignment of the *i*-th example with the k-dimensional vector **q**_*i*_, where *q_ij_* = 1 if the *i*-th example belongs to the *j*-th cluster and 0 otherwise.

We learn a representation of the patients to predict the cancer type of a given patient and the survival time. Every sample is associated with a class label that denotes the diagnosed cancer type of the patient *i, y_i_* ∈ {1, …, *m*}, where m is the number of cancer types. We denote the patient’s survival time with the *i*-th sample as *t_i_* and the patient’s survival status with *c_i_*. *c_i_* is 1 if the patient has passed away and 0 if it is censored. The cancer patient data, *D*, can be summarized as 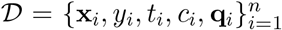. Here, the cluster membership vector **q** is unobserved.

#### 2.2.1 DeepCrossCancer Clustering Architecture

DeepCrossCancer’s clustering network consists of an input representation module, a classification module, a survival module, and a clustering module (Figure 1). While the classification module aims to categorize the patients into the correct cancer type, the survival module aims to predict the survival times of the patients accurately. The representation module takes the input, forms a nonlinear transformation of the data using multiple hidden layers, and projects it into a lower-dimensional space in the last hidden layer. This encoding layer is connected to the output layer for the classification and survival prediction. This encoding of the inputs is used in the clustering module. In this way, DeepCrossCancer clusters the patients with a learned representation of the inputs that can achieve classification and survival prediction. The network is trained to accomplish these tasks jointly, in which we provide the details in Section 2.2.3.

**Figure 1:**
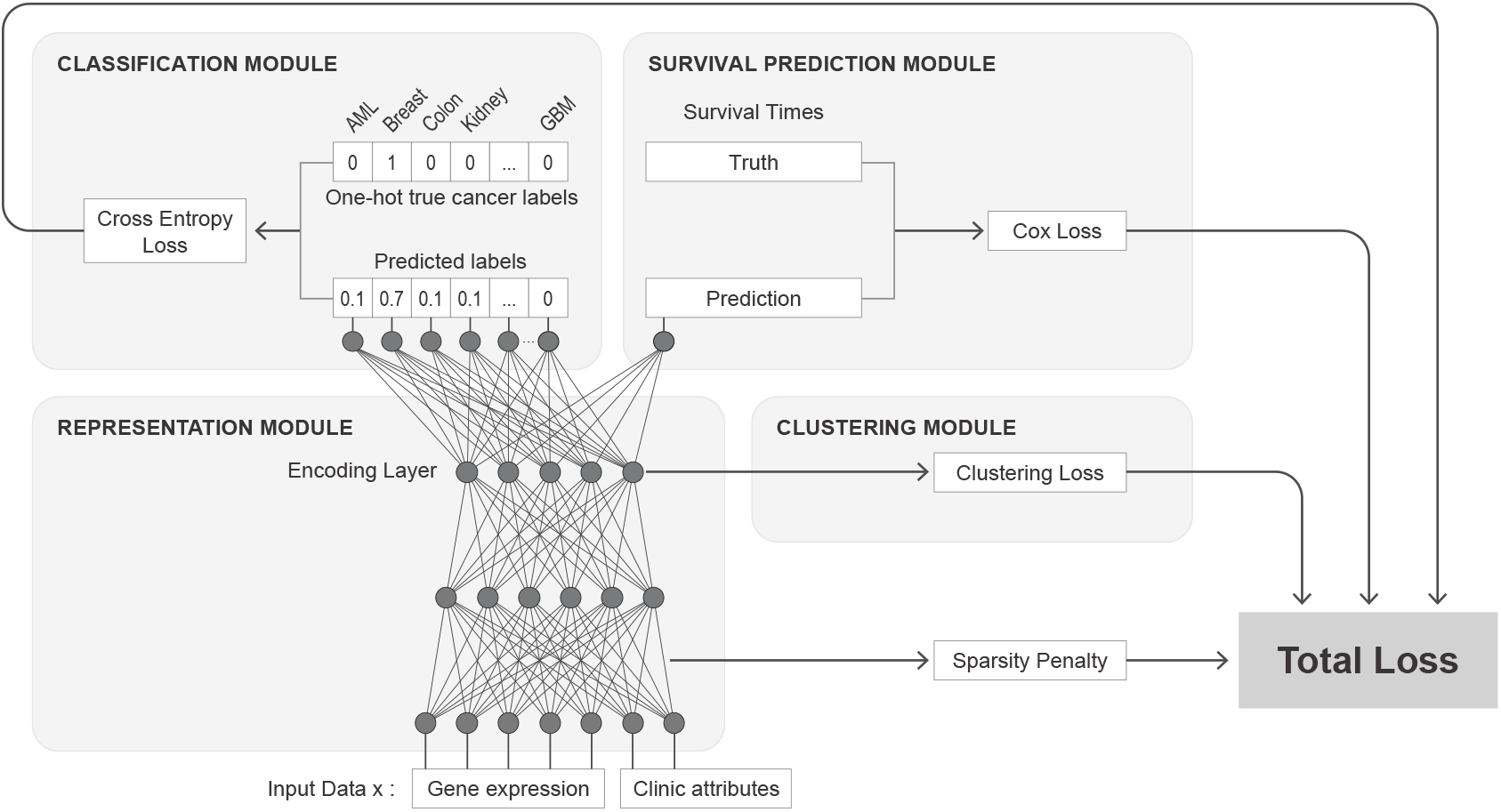
The network consists of four main components: representation, classification, survival, and clustering modules. The representation module applies a nonlinear transformation on the input data and maps them into a lower-dimensional representation on the encoding layer. The clustering module uses the representation provided in the encoding layer to group patients into *k* clusters.

The formulation and the architecture of the clustering network are built upon DeepType [25], which also uses representation, classification, and clustering modules. Different from DeepType, DeepCrossCancer’s clustering network contains an additional survival module. Note also that the classification modules in these two methods serve two different purposes. While DeepType’s classification module’s goal is responsible for classifying the patients of the same cancer type into prior known subtypes, in DeepCrossCancer, the classification module focuses on classifying patients from multiple cancer types into the diagnosed cancer types of the patient. The network used in DeepCrossCancer can be summarized as follows:

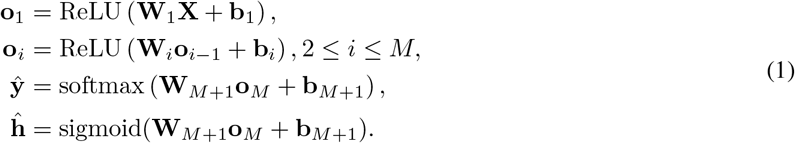

**W**_*i*_ denotes the weight matrix, **b**_*i*_ is the bias term vector and **o**_*i*_ is the output of the i-th layer. ŷ denotes the classification output and **h** is the survival output. Θ denotes the learnable network parameters Θ = (**W**, **b**). The RELU activation function [26] is used in the hidden layers; while softmax activation and sigmoid activation functions are used for the classification and survival layers, respectively. The network parameterized with Θ transforms inputs into a lower dimensional (*p* ≪ *d*) *latent* feature space *Z* in 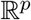 at the last hidden layer, *f_Θ_*: *X* → *Z*. Instead of clustering directly in the original data space 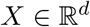, the clustering module uses this transformed representation of the inputs. The transformed data points, 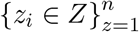 and the *k* cluster centers lie in 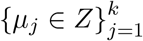 in this latent feature space *Z*.

#### 2.2.2 Loss Functions

The network is optimized jointly to achieve success in the three learning tasks. Supervised by the classification labels and the survival times, the network learns a representation that would lead to a latent space, *Z*, which will be useful in the unsupervised learning conducted for clustering. The network parameters are learned by minimizing the following objective function:

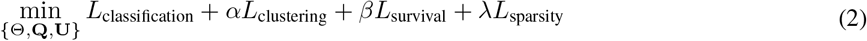

**U** is the centroid matrix. Each column represents a cluster center and is hidden. The **Q** is the cluster membership matrix, **Q** = [**q**^(1)^,…, **q**^(n)^], each row being one patient’s assignment vector. The parameter *λ* is the regularization parameter that controls the model sparsity, and *α* and *β* are parameters that adjust the importance assigned to the clustering and survival losses relative to the classification loss.

We use the cross-entropy loss to quantify the discrepancy between the correct cancer type of the patient and the predicted cancer types of the patient. As in [25], we impose an *ℓ_1_* regularization [27] on the weight matrix of the first layer to control the model complexity. The cross-entropy and sparsity losses are defined as:

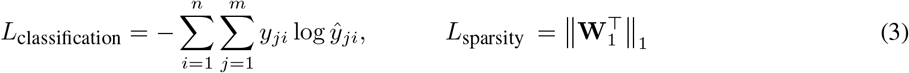

We use the k-means [28] loss that quantifies the tightness of the clusters around their centroids:

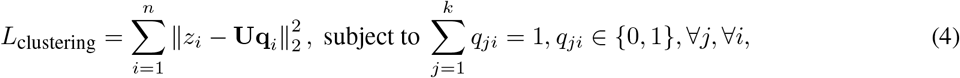

The survival module follows the Cox partial likelihood model [29].We use the Cox loss as defined in [30]:

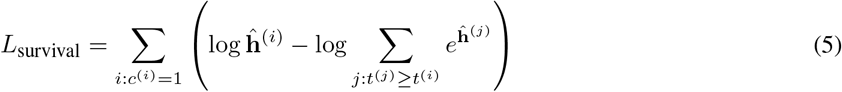

#### 2.2.3 Parameter and Hyper-parameter Optimization

The optimization problem should solve for **Θ**, the network parameters, and **U**, the centroids of the clusters, and **Q**, the assignment of the clusters simultaneously. Since they are coupled, as in DeepType, we employ an alternating minimization strategy. Initially, we ignore the clustering module by setting *α* to zero and pre-train the network to find an initial set of values for **Θ** and the parameters *β* and *λ*. We fix **Θ** and calculate the transformed points, *z_i_ ∀i*, and using standard k-means algorithm finds the clusters; thus, **Q** and **U**. In the next step, we use the **Q** and **U** found in the previous step to optimize for *θ*. We iterate these two steps alternatively until convergence. When training the network, we employ back-propagation by using the mini-batch based stochastic gradient descent method [31]. The hyperparameters associated with each loss, *α*, *β*, and *λ*, needs to be tuned as well. Since a grid search strategy for these three parameters is computationally expensive, we first optimize *β* and *λ* by setting *α* = 0; then we tune *α* for the best *β* and *λ* values. We use ten-fold stratified cross-validation on the training data. We use the random search with a probabilistic reduction optimization strategy provided in Talos optimization tool [32]. Detailed descriptions of these steps are provided in Algorithms 1, 2, 3 and Supplementary Text.

#### 2.2.4 Additional Evaluation Metrics

In addition to the losses defined for each task, we rely on different evaluation metrics for assessing the performance of the different components. We use the concordance index (C-index) to evaluate the survival module [33]. C-index calculates the fraction of patient pairs that are predicted to have the correct partial rank among all comparable pairs. To assess cluster quality, we use the silhouette score [34]. The silhouette score measures an object’s similarity to its cluster compared to other clusters. A value close to 1 indicates that the instance is assigned to its own cluster, whereas a value close to −1 means that it is closer to another cluster than its own. We use the silhouette score also to find the cross-cancer patients as detailed in the next section.

### 2.3 Step 2: Identifying Cross-Cancer Patients

We group the patients into clusters for a set of increasing values of *k*, we call this ordered set of *k* values *K*. Next, we compute a patient-by-patient similarity matrix. *(i,j)* entry of the matrix will indicate the fraction of times the patients co-clusters over the |K| many clusterings. To find similar patient pairs, we only consider those pairs where they always co-cluster. The reason for this choice is that we aim for a high precision than a high recall in detecting cross-cancer patients. Since there are no gold standard labels to evaluate precision/recall, we pick the highest possible threshold that would lead to high precision. This choice can be revisited in applying the model in different setups.

Once the similar pairs are identified, we find the cross-cancer patients by checking whether they are closer to their cancer type patients or the similar patients’ cluster. For this, we use the sign of the silhouette score. In the transformed space, *Z*, we calculated the silhouette score; in these calculations, the patients’ clusters are the diagnosed cancer types of patients. A positive silhouette score indicates that the patient is close to other patients diagnosed with same cancer, and such a patient can well be a representative member of that cancer type. On the other hand, the negative silhouette score indicates that this patient is closer to other cancer-type patients than patients with the same cancer. In summary, a patient who has a similarity score of 1 with another patient from another cancer type and with a negative silhouette score in all clustering models are deemed as a cross cancer patient. For a cross-cancer patient *i*, we will denote the set of patients to whom this patient is similar to with the patient set *S_i_.*

### 2.4 Dataset Processing

We use the TCGA (The Cancer Genome Atlas) patient data [35]. We obtain the processed gene expression data and clinical information of ten different types of cancer from [36]^1^. Patient somatic mutation information is obtained directly from the Broad Institute Firehose repository [37]. Patient copy number variation data were obtained from the Broad Institute TCGA GDAC Firehose repository by using the RTCGA-Toolbox, version 2.16.2 [38]. The following cancer types are covered in the dataset: acute myeloid leukemia (LAML), breast invasive carcinoma (BRCA), colon adenocarcinoma (COAD), kidney renal clear cell carcinoma (KIRC), liver hepatocellular carcinoma (LIHC), lung squamous cell carcinoma (LUSC), skin Cutaneous Melanoma (SKCM), ovarian serous cystadenocarcinoma (OV), sarcoma (SARC), and Glioblastoma multiforme (GBM). In the following analysis, we refer to these cancer types as AML, breast, colon, kidney, liver, lung, melanoma, ovarian, sarcoma, and GBM, respectively. In our analysis, we use only primary solid tumor samples. Since most samples are metastatic samples for melanoma cancer, and there are only 103 primary solid tumor samples, we decided to exclude melanoma and were left with nine cancer types. The number of patients ranges from 161 AML to 1211 breast cancer samples (Supplementary Table S2). The gene expression is quantified by the RNA-seq experiments and was processed by the RNA-Seq Analysis pipeline of TCGA. The gene expression values are normalized with RSEM count estimates and cover 20,531 genes. We obtain age, sex, and survival information from clinical annotation data provided by the TCGA. The survival time is the number of days to the last follow-up if the patient is alive and the number of days to death if the patient has passed away. We discretize age into eight bins. The discretized age bins are 0 – 20, 20 – 35, 35 – 45, 45 – 55, 55 – 65, 65 – 75, 75 – 85, and 85 —…. Patient somatic mutation information is obtained directly from the Broad Institute Firehose repository [37]. Patient copy number variation data from GISTIC2.0 (last analyze date 20160128) [39] were obtained from the Broad Institute TCGA GDAC Firehose repository by using the RTCGA-Toolbox, version 2.16.2 [38].

## 3 Results

### 3.1 Experimental Set-up

We use gene expression data of tumors biopsied from cancer patients and the clinical annotation data made available by the TCGA project (see Section 2.4 for details). We held out 20% of the data as test set, features are normalized into the range [0-1] on the training data. We design a five-layer neural network: an input layer, two hidden layers, a classification layer, and a survival prediction layer. The number of nodes for the input layer, hidden layers, classification layer, and survival layer is set to 20533, 32, 16, 9, and 1. Learning rate was 0.0001. The batch size and the number of epochs are set to 30 and 200, respectively, for the first part of the iterative optimization. For the second, step the learning rate was set to 0.05; the batch size and the number of epochs are 24, and 150 for training. For the clustering part, we train seven clustering models with seven different numbers of cluster size *k.* By using the method proposed in Section 2.2.3, the regularization parameter *λ*, and the trade-off parameters *α*, and *β* are optimized in each iteration. We use Adam [40] and SGD [31] optimizers in training the model.

### 3.2 Evaluating Clusters

We evaluate the performance of clustering for different values of *k* ∈ **K** = [10, 20,30,40, 50, 70,100]. The evaluation metrics for *k* are reported in Table 1. The high silhouette score, accuracy, and C-index demonstrate that the models are well trained. The optimal cluster number is 10 based on silhouette score; it includes all cancer types as separate clusters as expected and divides BRCA into two clusters. And not surprisingly the silhouette score degrades as the *k* increases. This is not a problem for our purposes as we are looking for patient pairs that remain to be in the same cluster even in that extreme clustering situations.

**Table 1:** The performance measures on the test data are reported with different numbers of clusters (k).

| Performance metrics/k | 10 | 20 | 30 | 40 | 50 | 70 | 100 |
| --- | --- | --- | --- | --- | --- | --- | --- |
| Accuracy | 0.97 | 0.98 | 0.97 | 0.98 | 0.98 | 0.97 | 0.98 |
| C-index | 0.69 | 0.70 | 0.73 | 0.69 | 0.72 | 0.71 | 0.72 |
| Silhouette score | 0.65 | 0.44 | 0.33 | 0.28 | 0.26 | 0.24 | 0.22 |

We compare DeepCrossCancer in terms of clustering performance with several other algorithms. The compared algorithms include the spectral clustering algorithm [41] with RBF kernel, k-means when features are included as is, k-means where an autoencoder processes the features. The parameters of these compared models are provided in Supplementary Text Section 5. Spectral clustering and k-means do poorly in terms of silhouette score compared to all variant of DeepCrossCancer (Figure 2).

**Figure 2:**
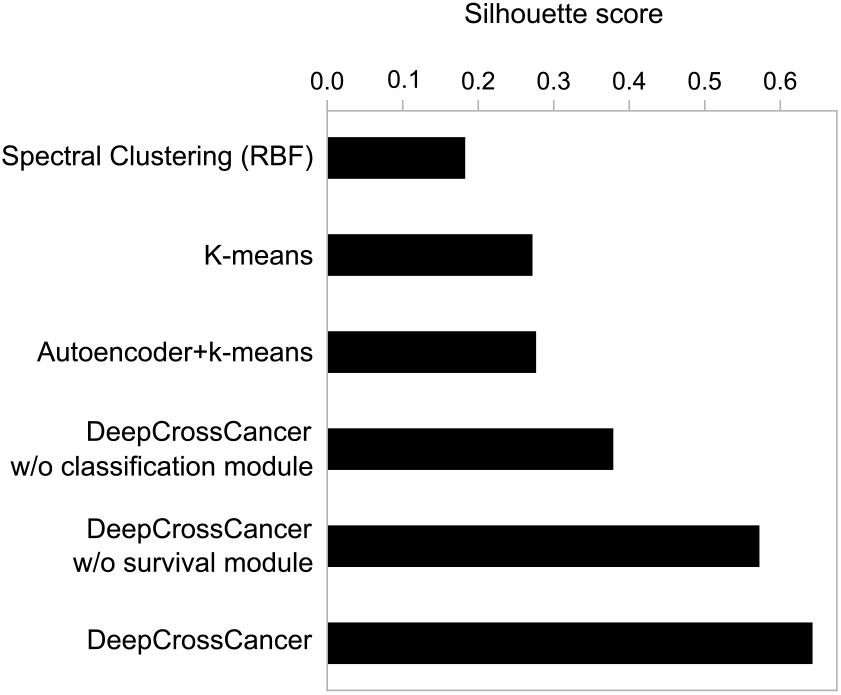
Comparison of silhouette scores of DeepCrossCancer with five other clustering algorithms for the number of clusters 10.

We also assessed whether supervision contributes to overall clustering success by training DeepCrossCancer without supervised parts. We obtained results for DeepCrossCancer without the classification module and without the survival module separately. The comparisons show that DeepCrossCancer results in best silhouette scores (Figure 2). Turning of the classification module degrades the performance most (41.05 %). Although the performance decrease for the survival module is not that large, its presence improves the performance by 10.9 %.

### 3.3 Identified Cross-Cancer Patients and their Similar Patient Set

To visualize clusters obtained, we apply t-SNE on the output of the encoder of the proposed model with *k* = 100 (Figure 3c). The figure is colored based on the diagnosed cancer types, and it shows that there are patients further away from their own cancer type members. For example, some breast, kidney, and liver cancer patients are closer to sarcoma patients. To find the patient pairs that are similar at all clusterings, we calculate a similarity score per patient pair based on how frequently the patient pair co-cluster (see Section 2.3). Figure 3a represents the heatmap of patients’ similarity scores.. Off-diagonal black points show similar patients across cancers. The distribution of similarity scores of patients is shown in Figure S2. We only consider patient pairs with a similarity score of 1 because these patients are grouped into the same cluster across all models with different numbers of clusters. Figure 3b shows similar patient pair distributions across different cancer types. Most similar patients are related to sarcoma.

**Figure 3:**
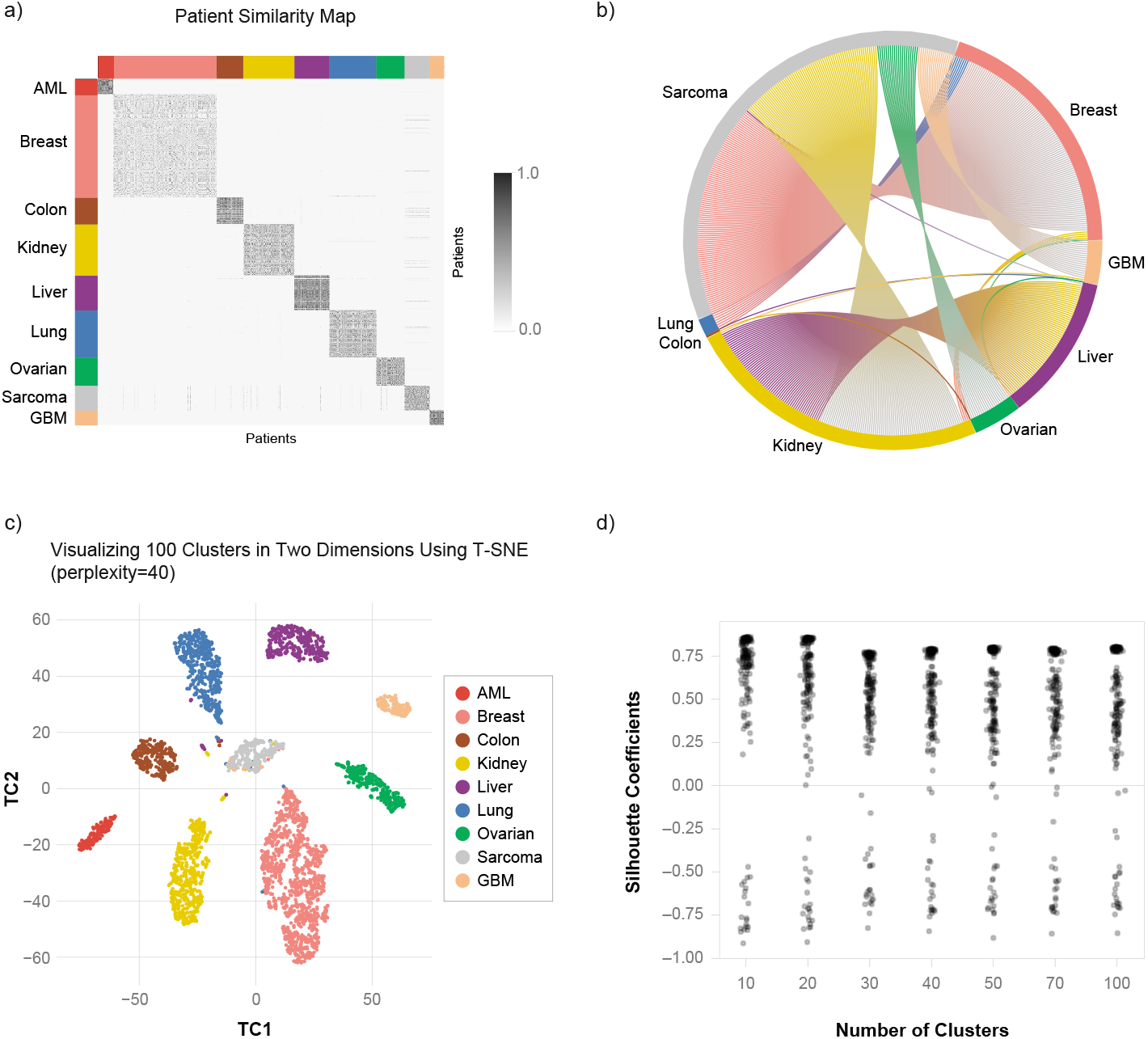
Cross-cancer patients are revealed in the different types of cancer. (a) The pairwise similarity of all patients is visualized by the heatmap. The similarity is based on how often the patients co-clusters. Off-diagonal black points represent similar patients across cancer. (b) Similar patients across cancers are shown by the chord diagram. (c) Example t-SNE plot for clustering with DeepCrossCancer with *k* = 100. The patients are colored by the actual cancer types. (d) The distribution of the silhouette coefficient of cross-cancer patients shown. Patients with a negative silhouette coefficient among similar patient pairs are the cross-cancer patients.

Of similar patient pairs, not all members are cross-cancer patients (Section 2.1 and Section 2.3). For example, as seen in the t-SNE graph (Figure 3c), one patient could be in the center of its cluster member; thus, the other patient, though, could be the outsider to its cancer type cluster, and then that patient will be the cross-cancer patient. To find such cross-cancer patients, we use silhouette coefficients. In these calculations, clusters are based on the actual cancer type classes, but the representation is obtained from the deep learning model’s encoding layer. Patients that always have a negative score at all clusterings are deemed cross cancer patients. 20 out of 176 patients were found (Figure 3d). We will refer to these 20 patients as the cross-cancer patients and are shown in Figure 4 as the center node with the patients that are found similar to. These patients appear in eight cancer types, except AML. None of the AML patients had a cross-cancer relationship. The reason might be that AML is a type of blood cancer, whereas the sample type of other cancers is a primary solid tumor. The list of patients is provided in Supplementary File 1. We present an analysis of cross cancer patients, along with those who exhibit similarities in the upcoming sections.

**Figure 4:**
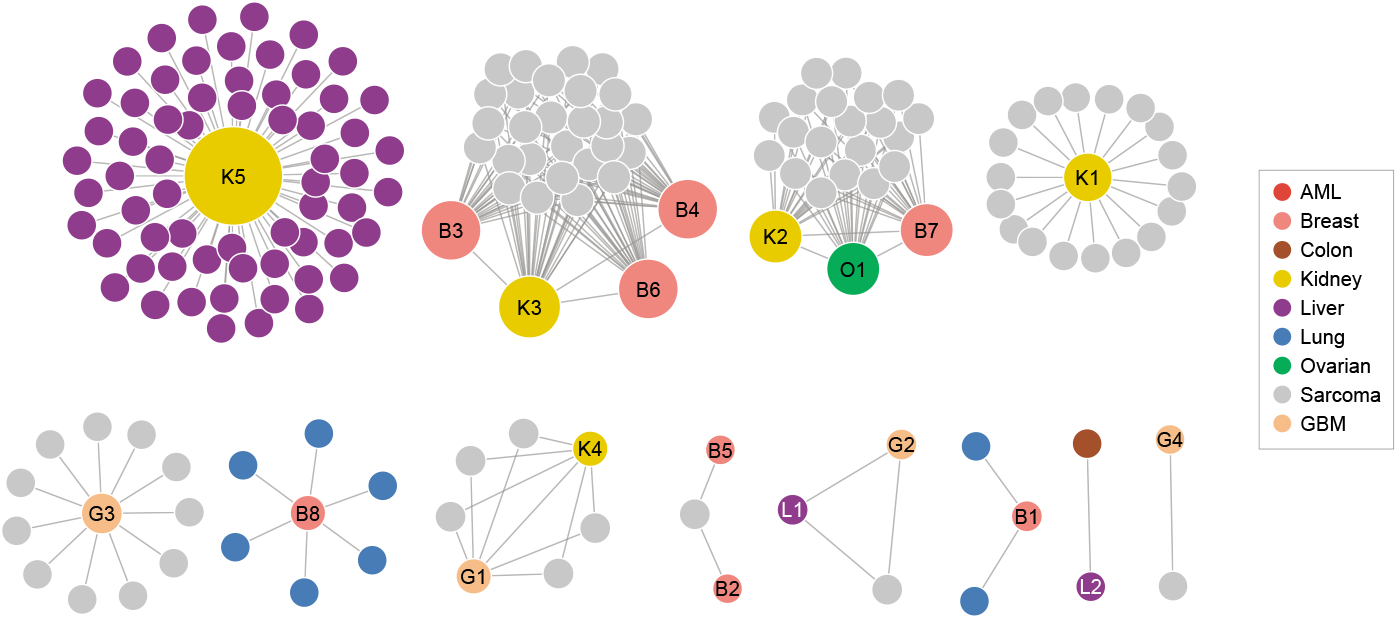
The network of cross-cancer patients. The relationship of patients across cancers is shown in the network. Cross-cancer patients are assigned an ID and are shown in the center of the network. The TCGA study abbreviations for cancer types in the legend are as follows: LAML, BRCA, COAD, KIRC, LIHC, LUSC, OV, SARC, and GBM. TCGA patient IDs of the patients are listed in Supplementary File 1.

Since there is no ground-truth list of cross-cancer patients, we applied the cross-cancer identification steps (Section 2.3) to the results of K-means and Spectral clustering. We repeatedly clustered, find the co-clustering pairs and among them identified the list of cross-cancer patients that these algorithm suggests. The numbers of cross-cancer patients found by these algorithms and DeepCrossCancer are shown in Supplementary Figure S3. K-means and spectral clustering suggest a very large number of cross-cancer patients. 14 out of 20 cross-cancer patients that found by DeepCrossCancer, are also found by K-means and Spectral clustering and 3 of them are found by K-means.

### 3.4 Shared Predictive Genes

For each cross-cancer patient identified (the center node in Figure 4), we have a set of patients that this patient is similar to (the connected nodes to the center node in Figure 4). First, we identify the predictive genes for each of the patients and check if these predictive genes are shared across the cross-cancer patient and its similar patient set. To quantify the importance of a gene in clustering for a patient, we use the Deep SHAP method, which relies on Shapley values to explain the contribution of an input feature [42]. We define 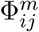 as the SHAP value for each patient *i*, and input feature *j* in model *m* of the |K| different models. In finding the SHAP values, we fit the DeepExplainer by specifying the clustering part. To find the genes that consistently emerge as the important features, we consider the genes in the top 1 percentile of all genes when ranked based on the SHAP values in all clusterings. The pseudo-code of the proposed procedure is in Algorithm 4. We next test whether the number of shared genes is statistically large.

We use a non-parametric permutation test for testing the null hypothesis that an observed statistic is significantly large. Let *i* denote a cross-cancer patient, and *S_i_* is the set of patients that this patient is similar to. *B* is the number of permutations, a test statistic *t* is calculated over patient *i*, and the patients in *S_i’_* drawn from the population in the cancer type of *i*. *S_i’_* patients are drawn *B* times and alternative test statistics (*t’*) is calculated over *i* and *S_i’_*. Based on the number of times *t* ≤ *t’* in *B* samplings (let it be *c* times), the empirical test statistic is calculated by the ratio *c/B*. Similar permutation tests are also used in the following parts.

In the analysis of SHAP genes, the test statistic is the number of common genes between the patient *i* and its similar patient set *S_i_*. The number of samplings B is set to 10,000. We repeat the method for each cross-cancer patient. We use Benjamini and Hochberg (B&H) correction for multiple hypothesis test correction [43]. For example, the kidney patient (K5) shares 13 common predictive genes with 63 liver patients in the first cross-cancer subnetwork. The number of common genes is always bigger than the number of common genes for the randomly selected 63 kidney patients (p-value ≤ 0.0001). This analysis is repeated for every cross-cancer patient; there are 8 cross-cancer patients sharing a significantly large number of genes with their similar patient set at significance level 0.05. The detailed list is provided in Supplementary File 2.

### 3.5 Gene Expression Analysis of the Cross-cancer Patient K5

To further understand the nature of the similarity between these cross-cancer patients, we conduct a gene expression analysis. We identify the genes for which the expression level is more typical for the cross-cancer type rather than the patients’ diagnosed cancer type. Here, we specifically focus on for K5 due to the large set of similar patients. We find genes, for which the expression level in K5 are not typical in kidney cancer but are more like the liver patients’. To this end, we calculate two z-scores. *z_i_*^(1)^ is the z-score of gene *i* for a particular cross-cancer patient, calculated by the sample mean and standard deviation of gene expression calculated over the cancer type of the cross-cancer patients; whereas *z*^(2)^ is calculated with the mean and standard deviation of the 63 liver patients to which K5 is found similar. We define Δ*z* = |*z*_2_| − |z_1_|. For a gene, if the patient is more similar to the cross-cancer type patients, Δ*z* will be negative. We find the genes Δ*z* is significantly large and negative using z-test and applied B&H correction [43], where the number of tests is the number of genes.

Significant genes at an FDR cut off of 0.1 for the cross-cancer patient (K5) are listed in Supplementary File 3. Figure 5a shows the top 15 genes, the expression values of kidney and liver patients are shown for the cross-cancer patient K5. The yellow-point represents K5, and purple-points represent liver patients that are similar to K5. For example, CYPY2A13 is over-expressed in K5, while this expression level is atypical for kidney it is typical for the 63 liver cancer patients to which K5 is similar. We also repeat the same analysis by controlling age and gender. To this end, we only take into account the patients with the same gender and the same age range as the cross-cancer patient. Gene expression profiles with the controlled gender and together with age for the cross-cancer patient are given in Supplementary Figures S4. When we control the age and sex, the number of significant genes is decreased; however, the top most significant genes are almost the same as the previous analysis results.

**Figure 5:**
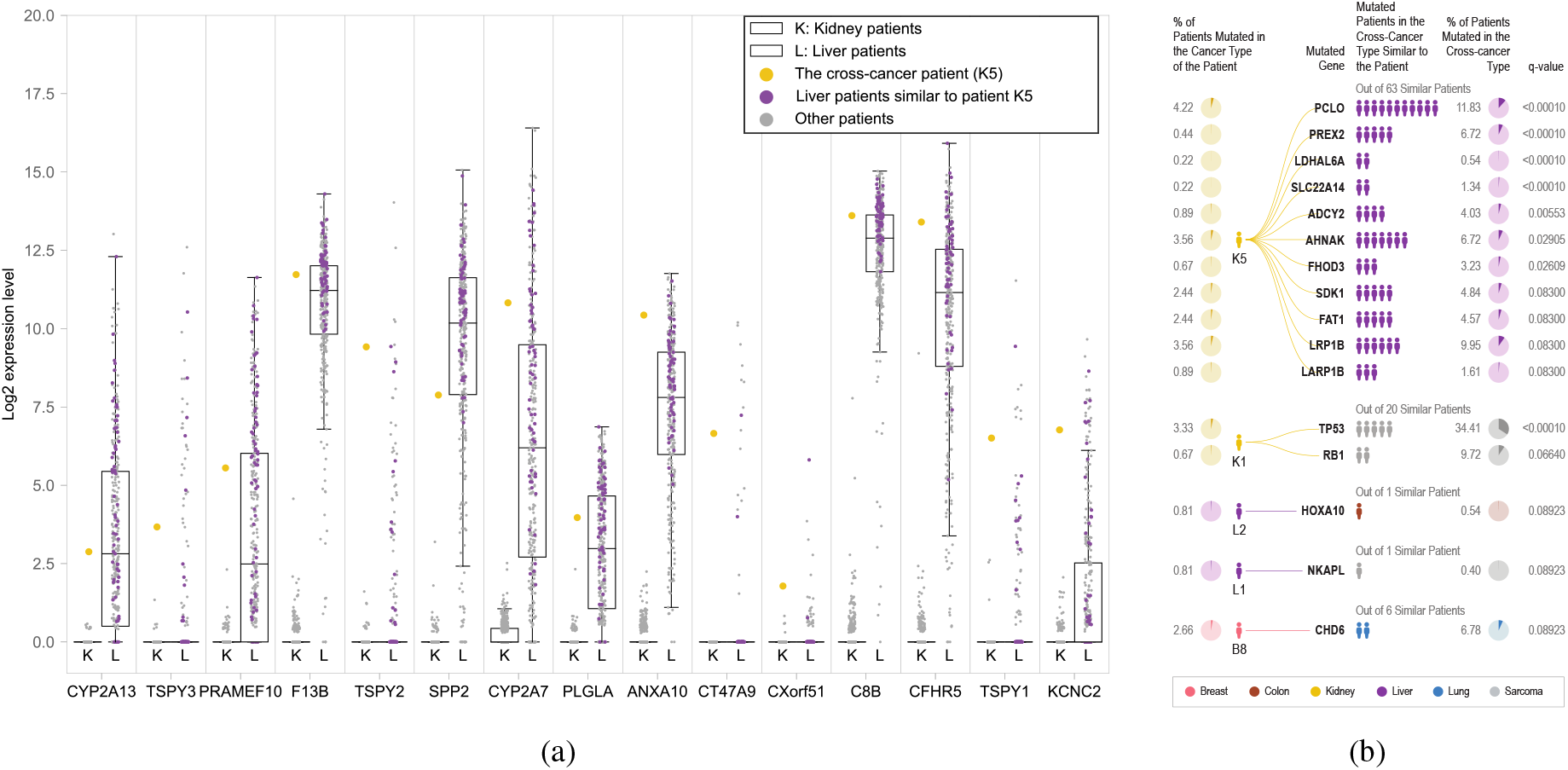
(a) The cross-cancer patient K5 is represented with yellow-point and liver patients that are similar to K5 are shown with purple points. The most 15 significant genes (q-value ≤ 6.83e-14) are listed in the figure. Others are in Supplementary File 3. (b) Five cross-cancer patients share a significant number of commonly mutated genes with similar patients. These mutated genes appeared with a 0.1 FDR threshold. Details of the figure are shown in Table S3.

Among the top ranked genes, there are members of cytochrome P450 superfamily of enzymes. The CYP enzymes complex units, particularly members of the CYP2A gene family, CYP2A13 and CYP2A7 have elevated gene expression levels unlike the kidney patients and similar to the liver patients. These genes which are known to be associated with drug metabolism.

### 3.6 Analysis of Shared Mutated Genes

We analyze the mutated genes of a cross-cancer patient and the patients that they are similar to. For each gene that is mutated in the cross-cancer patient, *i*, and in at least one of the similar patients set, we conduct a permutation test analysis to test if there are significantly many patients in the similar patient group, *S_i_*, that bears the same mutated gene. When conducting the test for gene *g* and for patient *i*, the test statistic *t* is the number of patients in *S_i_* for which *g* is mutated. The same statistic is calculated over random samplings of the cancer type of the cross-cancer patient. For example, when testing the significance of K5, 63 random patients are selected from the kidney; note that this is a more difficult test than selecting the random samples from the liver patients.

16 genes are found significantly shared among the cross-cancer patient it under the significance level with an FDR threshold of 0.1. These genes are *TP53, RB1, HOXA10, CHD6, NKAPL, PCLO, PREX2, LDHAL6A, SLC22A14, ADCY2, AHNAK, FHOD3, SDK1, FAT1, LRP1B,* and *LARP1B* and are are provided in Figure 5b with the associated cross-cancer patients the overall mutational frequency of these genes in the cross-cancer patient’s cancer type and the cross-cancer type. There are interesting findings. For example, while TP53 is the most frequently mutated gene in most human cancers [44], TP53 mutation is observed in only 3.33% of all kidney cancer patients. One of the cross-cancer patients (K1) is found to be similar to two sarcoma patients and TP53 is found to be one of the significantly shared genes for this relationship. Interestingly, 34.4% of sarcoma patients carry a TP53 mutation. Gurova et al. found that renal cell carcinoma (RCC) rarely acquires mutations in the p53 tumor suppressor gene, and the p53 signal is suppressed by another mechanism in this tumor type [45]. A similar case is observed RB1 gene, while it is not frequently mutated in the kidney cohort, it is frequently mutated in the sarcoma and these kidney patients are found to be similar to sarcoma patients. In a new approach called MutPannig, TP53 and RB1 have also found to be the most important mutated genes as driver mutations in sarcoma patients [46].

### 3.7 Analysis of Shared Copy Number Alterations

Next, we analyzed significantly shared copy number alterations (CNA) between the cross-cancer patients and their similar patient sets. Since the number of copy number altered genes are large, we limit our analysis to cytobands with an alteration that is observed both in the cross-cancer patient, *i*, and in at least 70% of their similar patient set, *S_i_*. We analyzed the amplification and deletion events separately. For patient *i*, the test statistic *t* is the number of patients in *S_i_* with deletion (or amplification) in cytoband *c*, and alternative test statistic *t*’ is calculated over the randomly selected patients from the cancer type of the cross-cancer patient. For *B* = 1000, we repeated the method for each common thresholded cytoband.

55 cytobands in amplification events rejects the null hypothesis test with FDR threshold of 0.1 (q-value ≤ 0.066). Four cross-cancer patients, G3, G1, K4, and B8, have shown significant amplification events shared with their similar patients. The most frequent chromosomal arm alterations included copy number gains in 1q and 1p of the cross-cancer patient G3, in 17p and Xp of G1, in 4p, 17p and Xp of K4, and in 3q of B8. Since G1 and K4 are connected in the same subnetwork, they show the common frequent copy number gains in 17p and Xp. The significantly amplified cytobands are listed in Supplementary File 6. 119 cytobands with deletions pass the significance test with an FDR threshold of 0.1 (q-value ≤ 0.094). Six cross-cancer patients - G1, K4, K1, K2, B7, and B8 - have shown significant deletion events shared with their cross-cancer similar patient set. The significant chromosomal losses are in 16q and 17p of the cross-cancer patient G1, in 1q, 2q, 4q, 10p, 10q, 13q, 17p, 18q, and Xq of the patient K4, in 13q of K1, in 13q and 16q of K2, in 13q of B7, in 4p, 4q, and 9p of B8. G1 and K4 have the common copy number losses on the chromosomal arm 17p, and K2 and B7 have the common losses on the chromosomal arm 13q. These significantly deleted cytobands are listed in Supplementary File 7.

## 4 Conclusion and Future Work

In this work, DeepCrossCancer, for discovering cross-cancer patients using patient molecular profiles and clinical information. Using DeepCrossCancer, we find 20 cross-cancer patients that cross boundaries in nine cancers. Analysis of these patients in light of their predictive genes and other genomic information led to interesting findings. For example, for a kidney diagnosed patient, whom we find similar to liver cancer patients, the shared common genes include genes reported to be prognostic markers for liver cancer. We also identified a set of loci and genes that are mutated or copy number altered in the cross-cancer patients. The study presents new opportunities to treat different cancer patients who share transcriptomic similarities but respond poorly to tissue-specific treatments. Transferring clinical treatment strategies from one patient to another patient could be critical for some patients. The identified common alterations could be further investigated experimentally to decipher the common molecular mechanisms shared across these patients.

The reasons why each of these cross-cancer patients similar needs further investigation. The underlying cause could be the exposure to common carcinogens or shared genomic architecture that predisposes them to cancer in the same way. There can also be shared unknown factors; for example, the same specific drug intake or radiation therapy may affect similarities in their transcriptomes. Since we do not have access to medication history, we cannot correct for such hidden factors. However, the fact that these patients also share other genomic alterations supports the case that the reason is not such a common therapy history.

The proposed method also has its limitations. Since deep learning models require large training set sizes, DeepCrossCancer is not applicable to small patient cohorts. There are also several routes for further investigation. As input to clustering, we used the transcriptome data and used the mutation and copy number to analyze the discovered cross-cancer patients. Instead, the framework can be extended to a multi-modal framework with other types of omic characterization of the patients, such as methylation, mutation, and copy number variation. Another research direction could be the integration of the functional gene sets, and interaction networks to learn a better representation of the patients. We also plan to extend this work to find similar cancers in patients from 33 cancer types made available in the TCGA project. The information used in the different modules can be expanded. In the classification module, the class labels could be enriched with the known subtype information of the cancer types. In the survival module, other clinical information such as the stage can be incorporated. Finally, although DeepCrossCancer is designed for identifying cross-cancer patients, the framework can be extended to other diseases such as neurological diseases where common genomic alterations are reported.

## Footnotes

1 http://acgt.cs.tau.ac.il/multi_omic_benchmark/download.html

