## Supplementary for "Identifying Cross-Cancer Similar Patients via a Semi-Supervised Deep Clustering Approach": SupplementaryText.pdf

### Supplementary Tables and Figures

### 1 Notation

| Symbol | Explanation |
|---|---|
| $X$ | input space $\in \mathbb{R}^d$ |
| $Z$ | latent space derived from the last hidden layer of the network input space $\in \mathbb{R}^p$, where $p << d$. |
| $\mathbf{x}$ | input feature vector for patient $\in X$ |
| $\mathbf{z}$ | transformed feature vector for patient $\in Z$ |
| $n$ | number of patients |
| $y$ | class label representing the patient's diagnosed cancer type |
| $\hat{y}$ | predicted class label |
| $m$ | number of cancer types, number of classes for the classification task |
| $t$ | survival time |
| $\hat{h}$ | predicted survival time |
| $c$ | censorship status of the survival time |
| $\mathbf{W}$ | network weight matrix |
| $\mathbf{b}$ | network bias term |
| $\mathbf{o}$ | network output layer |
| $M$ | number of layers in the network |
| $\mathbf{\Theta}$ | network trainable parameters $(\mathbf{W}, \mathbf{b})$ |
| $\mathbf{U}$ | cluster center matrix, where each column is a cluster center |
| $\mathbf{u}$ | cluster center for a cluster $\in Z$ |
| $\mathbf{Q}$ | cluster assignment matrix, where each row is a 0-1 assignment vector for a patient |
| $\mathbf{q}$ | cluster assignment vector. For example, for patient $i$, $q_{ji} = 1$ if the $i$-th patient belongs to the $j$-th cluster and 0 otherwise. |
| $\alpha$ | trade-off parameter for clustering loss |
| $\beta$ | trade-off parameter for survival prediction |
| $\lambda$ | regularization parameter |
| $k$ | number of clusters |
| $\mathbf{K}$ | the set of $k$ values used for clustering |
| $\Phi$ | Shap value |

Table S1: The notation used throughout the article.

### 2 Parameter and Hyperparameter Optimization

For each $\beta$ value, we fix $\beta$ and find the best $\lambda$ value in each of the cycles of 10-fold cross-validation. For each fold, we pick the best $\lambda$ by using the Talos optimization tool [1]. We use the random search with a probabilistic reduction optimization strategy provided in Talos. The strategy uses a probabilistic method to remove poorly performing parameter configurations from the search space by quantifying the decline in the specified reduction metric. We choose the reduction metric as the concordance index of survival time prediction. We obtain the average of the best $\lambda$ values for each fold for a set $\beta_l$ value, and we refer to this as $\lambda_l^{\text{avg}}$ in Algorithm 1. In this 10-fold CV procedure to optimize $\lambda$, we also obtain the average classification error $e_l^{\text{avg}}$ and the associated standard deviation of over the 10-folds $\sigma_l$ (Line 6 in Algorithm 1). Using the one-standard-error rule [2] the best $\lambda_{l^*}$ value is picked for the $\beta_l$ value. This procedure is repeated for each possible value of $\beta_l \in T = \{\beta_1, ..., \beta_L\}$. Next, we choose the optimal pair, $(\beta^*, \lambda^*)$ using the one-standard-error (Steps 10-13 in Algorithm 1).

Once the optimal $\beta$ and $\lambda$ parameters are obtained, the deep learning model is pre-trained with these values and $\alpha = 0$. The pre-trained model $m$-th layer is used to transform the feature matrix $\mathbf{X}$ to $\mathbf{Z}$ and this is input to k-means algorithm to get the cluster centers, $\mathbf{U}$ and the cluster assignments $\mathbf{Q}$. Secondly, we obtain the optimal $\beta$ and $\lambda$ and train the entire model to find the optimal $\alpha$ for each number of clusters. Again by applying the one-standard-error rule [2], we choose the optimal $\alpha$ values for each number of clusters. The pseudo-code of the proposed procedure is given in Algorithms 1, 2, and 3, and performs well in our numerical experience as shown in Figure S1.

**Algorithm 1** Hyper-parameter optimization $(\mathcal{D}_{tr}, \mathbf{X}, \mathbf{Z}, A, B, T, k)$

**Input:** Training data $\mathcal{D}_{tr} = \{\mathbf{x}_i, y_i, t_i, c_i\}_{i=1}^{n_{tr}}$ ($n_{tr}$ = the size of training data), $\mathbf{X}$, feature matrix, where $i$-th row is patient $i$'s feature vector, $\mathbf{Z}$, transformed feature matrix at the $m$-th layer of the network, $A = \{\alpha_1, ..., \alpha_J\}$, $B = \{\lambda_1, ..., \lambda_L\}$, $T = \{\beta_1, ..., \beta_L\}$, number of clusters $k$ .

**Output:** Optimized parameters $\alpha^*, \lambda^*, \beta^*$.

——————— **Optimize** $\beta, \lambda$ ———————
$\alpha \leftarrow 0$;
$E \leftarrow \emptyset$; // The set of average errors and standard deviations for each $\beta$ in $B$
**for** $l = 1$ to $L$ **do**
  $\beta = \beta_l$;
  $(e_l^{\text{avg}}, \sigma_l, \lambda_l^{\text{avg}}) =$ **`OptimizeLambdawithTalosCV`**$(\mathcal{D}_{tr}, B, \beta, \alpha)$; // 10-fold
  $E = E \cup \{(e_l^{\text{avg}}, \sigma_l)\}$;
**end for**
————— **Apply one-standard error rule** —————
Find the minimum avg classification error $e_0$ and one standard error $\sigma_0$ in $E$;
$l^* = \arg\max_{1 \le l \le L} l$, subject to $e_l^{\text{avg}} \le e_0 + \sigma_0$; // one-standard-error rule [2]
$\beta^* = \beta_{l^*}$;
$\lambda^* = \lambda_{l^*}^{\text{avg}}$;
————————————————
$f_\Theta =$ **`TrainNetwork`** $(D_{tr}; \lambda^*, \beta^*, \alpha)$;
$\mathbf{Z} = f_\Theta(\mathbf{X})$;
$(\mathbf{Q}_0, \mathbf{U}_0) =$ **`k-means`** $(\mathbf{Z}, k)$ ;
$\alpha^* =$ **`OptimizeAlphawithCV`**$(\mathcal{D}_{tr}, \mathbf{Q}_0, \mathbf{U}_0, A, \beta^*, \lambda^*, k)$;
**return** $(\lambda^*, \beta^*, \alpha^*)$

**Algorithm 2** **`OptimizeLambdawithTalosCV`**

**Input:** $\mathcal{D}_{tr}$, $B = \{\lambda_1, ..., \lambda_L\}$, $\beta$, $\alpha = 0$.

**Output:** Average classification error $e^{\text{avg}}$, standard deviation of classification errors $\sigma$, average of optimal $\lambda$ values $\lambda^{\text{avg}}$.

Randomly partition $\mathcal{D}_{tr}$ into ten folds;
**for** i= 1 **to** 10 **do**
  $(e_i, \lambda_i)$ = `OptimizeLambdawithTalos` $(\mathcal{D}_{tr}^{(i)}, B, \beta, \alpha)$;
  // gets optimal $\lambda_i$ for fold $i$ with Talos [1] and the associated error.
**end for**
Compute average classification error $e^{\text{avg}}$;
Compute standard deviation of classification errors $\sigma$;
Compute average of optimal lambda values $\lambda^{\text{avg}}$;
**return** $(e^{\text{avg}}, \sigma, \lambda^{\text{avg}})$

**Algorithm 3** `OptimizeAlphawithCV`

**Input:** $\mathcal{D}_{tr}$, $A = \{\alpha_1, ..., \alpha_J\}$, number of clusters $k$, $\beta^*$ best $\beta$ value, $\lambda^*$ best $\lambda$ value, $\mathbf{Q}_O$ cluster assignments obtained with the pretrained model, $\mathbf{U}_0$ cluster centroids obtained with the pretrained model.

**Output:** Best parameter $\alpha^*$.

**for** $j = 1$ to $J$ **do**
 $\alpha = \alpha_j$;
 $(e_j^{\text{avg}}, \sigma_j) = \texttt{10foldCV}(\mathbf{U}_0, \mathbf{Q}_0, \alpha, \lambda^*, \beta^*, )$;
 // $e_j^{\text{avg}}$ the average classification error over ten folds.
 // $\sigma_j$ the standard deviation of the ten folds.
**end for**
$j^* = \arg\max_{1 \leq j \leq J} j$, subject to $e_j^{\text{avg}} \leq e_0 + \sigma_0$; // one-standard-error rule
$\alpha^* = \alpha_{j^*}$;
**return** $\alpha*$

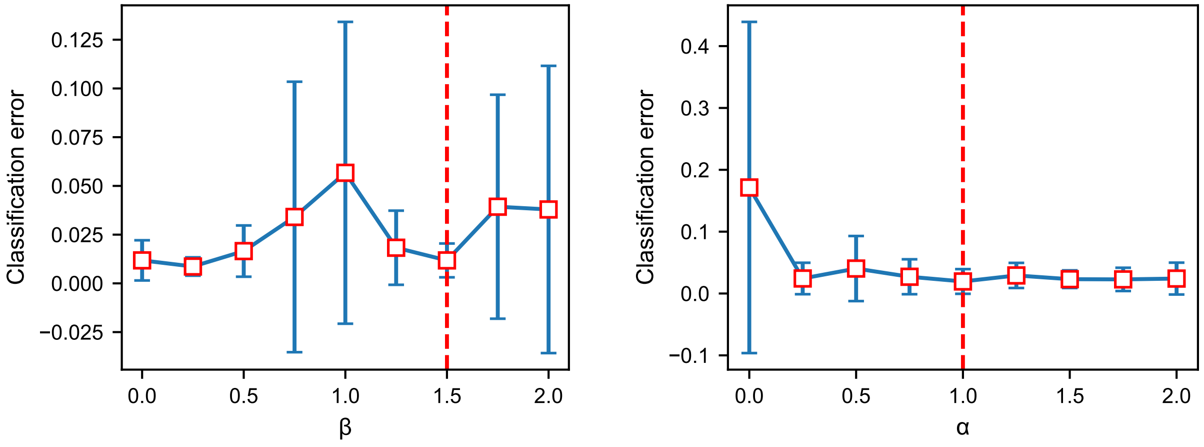


(a) (b)

Figure S1: Hyper-parameter optimization. (a) The optimal value of $\lambda$ is found to be 0.00056 by Algorithms 1 and 2. (a) shows the average classification error and the standard error over ten-CV folds. The optimal $\beta$ value is marked with the dashed red vertical line. (b) Example graph for the hyper-parameter optimization when $k = 10$. (b) shows the optimal value for $\alpha$ (see Algorithm 3).

### 3 Number of Patients

| Sample Type | AML | Breast | Colon | Kidney | Liver | Lung | Ovarian | Sarcoma | GBM |
|---|---|---|---|---|---|---|---|---|---|
| Primary Solid Tumor | 0 | 1077 | 278 | 537 | 367 | 489 | 294 | 258 | 151 |
| Recurrent Solid Tumor | 0 | 0 | 1 | 0 | 2 | 0 | 4 | 3 | 13 |
| Primary Blood Derived | 161 | 0 | 0 | 0 | 0 | 0 | 0 | 0 | 0 |
| Additional-New Primary | 0 | 0 | 0 | 1 | 0 | 0 | 0 | 0 | 0 |
| Metastatic | 0 | 7 | 1 | 0 | 0 | 0 | 0 | 1 | 0 |
| Additional Metastatic | 0 | 0 | 0 | 0 | 0 | 0 | 0 | 0 | 0 |
| Solid Tissue Normal | 0 | 111 | 40 | 72 | 48 | 51 | 0 | 2 | 0 |

Table S2: The number of cancer patients with sample types as in the dataset obtained from [3]. We use only primary solid tumor data.

### 4 Similarity Scores Distribution

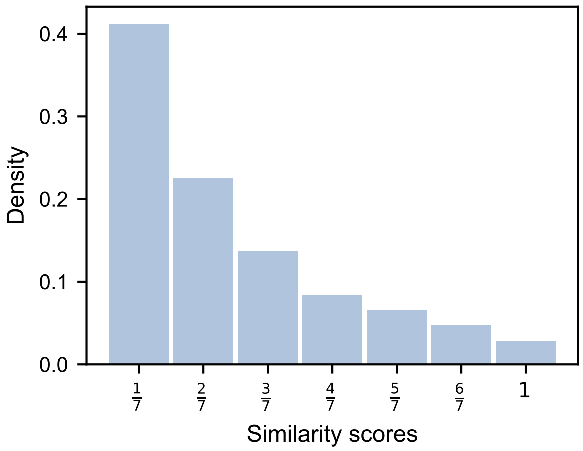


Figure S2: Similarity score is calculated for each patient pair as the fraction of frequency of co-clustering over multiple runs of clustering.

### 5 Parameter Settings for Different Clustering Models

We compare DeepCrossCancer with other clustering algorithms. One of them is spectral clustering. For spectral clustering, the affinity matrix is constructed using a radial basis function (RBF) kernel and the kernel coefficient for RBF (gamma) is 1. We also apply k-means on a basic autoencoder. The autoencoder is composed of the encoder, decoder, and bottleneck layer. For the autoencoder, we design a seven-layer neural network: an input layer, two hidden layers, a bottleneck layer, two hidden layers, and an output layer. The number of nodes for the input layer, hidden layers, and bottleneck layer are set to 20533, 32, 16, and 8, respectively. We also add $\ell_1$ regularization loss with the regularization parameter ($\lambda$) 1e-7 on the first and last hidden layers. The learning rate, the batch size, and the number of epochs are set to 0.001, 30, and 300. We use the mean squared error as a loss function and Adam as optimizer. We have validated the model, and overfitting and underfitting were not observed. Finally, we apply k-means on the bottleneck layer. For the other comparison models (DeepCrossCancer without supervised parts), we set the loss coefficient of supervised modules to zero.

#### 6 Overlapping of Cross-cancer Patients found by DeepCrossCancer and Other Clustering Algorithms

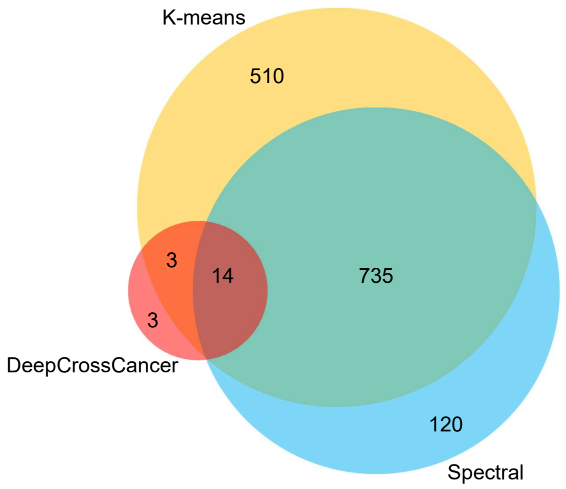


Figure S3: The Venn diagram shows the overlap of the cross-cancer patients for different algorithms.

#### 7 Shared Predictive Genes

**Algorithm 4** Getting Top Features with Deep SHAP

**Input:** List of number of clusters $\mathbf{K}$, the number of samples $n$, the set of similar patients $S^{(i)} = \{S_1^{(i)}, ..., S_s^{(i)}\}$ to the cross-cancer patient $i$, and $s$ is the number of similar patients.

**Output:** Common top feature list $P^{(i)}$ within the top $1\%$ between the cross-cancer patient $i$ and patients similar to the patient $i$.

**for** $k$ in $K$ **do**
  Load the trained model with the number of clusters $k$;
  Specify clustering part of the model $m$;
  Get SHAP values $\Phi^m$ with Deep SHAP;
  **for** $l = 0$ to $n$ **do**
    Take absolute values of $\Phi_l^m$;
    Get top features $P_l^m$ whose SHAP values within the top $1\%$;
  **end for**
**end for**
**for** $l = 0$ to $n$ **do**
  $P_l = \bigcap_{m=1}^{M} P_l^m$;
**end for**
Common top features between the cross-cancer patient $i$ and similar patients $S^{(i)}$:
$P^{(i)} = \bigcap_{l=S_1^{(i)}}^{S_s^{(i)}} P_l$; // Repeat for each cross-cancer patient $i$.

### 8 Gene Expression Analysis of the Cross-cancer Patient K5

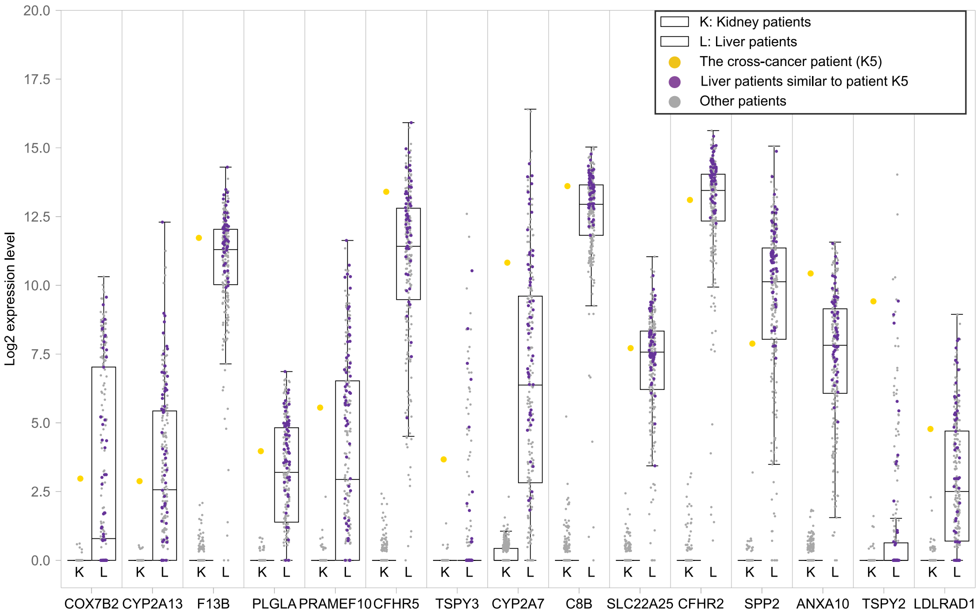


(a)

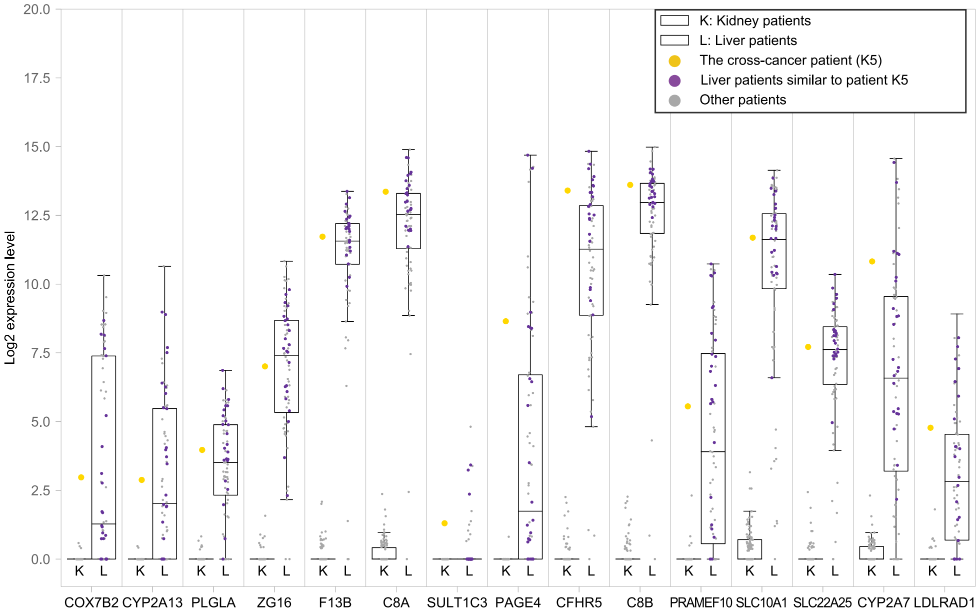


(b)

Figure S4: Gene expression profiles of subset of kidney (KIRC) and liver (LIHC) patients based on gender and age.(a) As a result of testing with gender subset on liver patients that are similar to K5 in Section 3.5, the most 15 significant genes (q-value $\leq 3.59$e-10) were represented. Details of the figure are shown in Supplementary File 4. (b) The test was done with liver patients who are similar to K5 in the same age and gender subset and the most 15 significant genes (q-value $\leq 5.04$e-3) were represented. Details of the figure are shown in Supplementary File 5.

### 9 Significantly Shared Mutated Genes

| TCGA Patient ID | Network Patient ID | Cross-cancer Type (Number of Patients Similar to the Patient) | Mutated Gene | % of Patients Mutated in the Cancer Type of the Patient | % of Patients Mutated in the Cross-cancer Type | Number of Mutated Patients in the Cross-cancer Type Similar to the Patient | B&H Adjusted P-value | Mutation Type in the Patient | Mutation Type in Patients of the Cross-cancer Type (Number of Patients in the Mutation Type) |
|---|---|---|---|---|---|---|---|---|---|
| TCGA-BP-4770 | K1 | Sarcoma (20) | TP53 | 3.33 | 34.41 | 5 | <0.00010 | Splice Site | Missense Mutation (4)<br>Frame Shift Del (1) |
| | | | RB1 | 0.67 | 9.72 | 2 | 0.066 | Splice Site | Splice Site (1)<br>Frame Shift Del (1) |
| TCGA-CC-5260 | L2 | Colon (1) | HOXA10 | 0.81 | 0.54 | 1 | 0.089 | Frame Shift Ins | Missense Mutation |
| TCGA-E9-A5FL | B8 | Lung (6) | CHD6 | 2.66 | 6.78 | 2 | 0.089 | Missense Mutation | Silent |
| TCGA-CC-A7IJ | L1 | Sarcoma (1) | NKAPL | 0.81 | 0.4 | 1 | 0.089 | Silent | Missense Mutation |
| TCGA-AS-3777 | K5 | Liver (63) | PCLO | 4.22 | 11.83 | 11 | <0.00010 | Missense Mutation | Missense Mutation (9)<br>Silent (2) |
| | | | PREX2 | 0.44 | 6.72 | 5 | <0.00010 | Splice Site | Missense Mutation (4)<br>Silent (1) |
| | | | LDHAL6A | 0.22 | 0.54 | 2 | <0.00010 | Silent | Missense Mutation (1)<br>Splice Site (1) |
| | | | SLC22A14 | 0.22 | 1.34 | 2 | <0.00010 | Missense Mutation | Missense Mutation (1)<br>Silent (1) |
| | | | ADCY2 | 0.89 | 4.03 | 4 | 0.006 | Missense Mutation | Missense Mutation (3)<br>In Frame Ins (1) |
| | | | AHNAK | 3.56 | 6.72 | 7 | 0.029 | Missense Mutation | Missense Mutation (3)<br>Silent (3)<br>Frame Shift Del (1) |
| | | | FHOD3 | 0.67 | 3.23 | 3 | 0.026 | Missense Mutation | Missense Mutation (1)<br>Silent (1)<br>Nonsense Mutation (1) |
| | | | SDK1 | 2.44 | 4.84 | 5 | 0.083 | Missense Mutation | Missense Mutation (4)<br>Splice Site (1) |
| | | | FAT1 | 2.44 | 4.57 | 5 | 0.083 | Missense Mutation | Missense Mutation (3)<br>Silent (1)<br>Frame Shift Del (1) |
| | | | LRP1B | 3.56 | 9.95 | 6 | 0.083 | Missense Mutation | Missense Mutation (4)<br>Splice Site (1)<br>Silent (1) |
| | | | LARP1B | 0.89 | 1.61 | 3 | 0.083 | Silent | Missense Mutation (2)<br>Nonsense Mutation (1) |

Table S3: The significance of commonly mutated genes was tested by a permutation test with B&H correction. Four cross-cancer patients show common genes that have been mutated significantly with patients similar to them.
